# Palbociclib-mediated cell cycle arrest can occur in the absence of the CDK inhibitors p21 and p27

**DOI:** 10.1101/2021.05.06.442976

**Authors:** Betheney R. Pennycook, Alexis R. Barr

**Affiliations:** Institute of Clinical Sciences, Faculty of Medicine, Imperial College London, Du Cane Road, London W12 0NN, UK; MRC London Institute of Medical Sciences, Imperial College London, Du Cane Road, London W12 0NN, UK

## Abstract

The use of CDK4/6 inhibitors in the treatment of a wide range of cancers is an area of ongoing investigation. Despite their increasing clinical use, there is limited understanding of the determinants of sensitivity and resistance to these drugs. Recent data has cast doubt on how CDK4/6 inhibitors arrest proliferation, provoking renewed interest in the role(s) of CDK4/6 in driving cell proliferation. As the use of CDK4/6 inhibitors in cancer therapies becomes more prominent, an understanding of their effect on the cell cycle becomes more urgent. Here, we investigate the mechanism of action of CDK4/6 inhibitors in promoting cell cycle arrest. Two main models explain how CDK4/6 inhibitors cause G1 cell cycle arrest, which differ in their dependence on the CDK inhibitor proteins p21 and p27. We have used live and fixed single-cell quantitative imaging, with inducible degradation systems, to address the roles of p21 and p27 in the mechanism of action of CDK4/6 inhibitors. We find that CDK4/6 inhibitors can initiate and maintain a cell cycle arrest without p21 or p27. This work clarifies our current understanding of the mechanism of action of CDK4/6 inhibitors and has implications for cancer treatment and patient stratification.

## Introduction

CDK4/6 inhibitors have garnered interest as cancer treatments due to their efficiency in inhibiting cell proliferation. Three small-molecule CDK4/6 inhibitors (Palbociclib, Abemabiclib and Ribociclib) are clinically approved for the treatment of metastatic ER+/HER2- breast cancer, and their use in the treatment of other cancers is an area of active investigation (Álvarez-Fernández & Malumbres, 2020; Dickler et al., 2017; Finn et al., 2016; Fry et al., 2004; Gelbert et al., 2014; Hortobagyi et al., 2016; Rader et al., 2013; Sledge et al., 2017; Tripathy et al., 2017). However, not all patients respond to these drugs and it is unclear why. Understanding more about the mechanism of action of CDK4/6 inhibitors, and how they inhibit cell proliferation, will help to stratify patients for treatment based on biomarkers (Álvarez-Fernández & Malumbres, 2020; Spring et al., 2020).

While the premise for the clinical use of CDK4/6 inhibitors is based on a “canonical” model of CDK4/6 activity, recent work has highlighted gaps in our understanding of the role of CDK4/6 in cell cycle entry (Pennycook & Barr, 2020; Hume et al., 2020; Rubin et al., 2020). In this canonical model, Cyclin D:CDK4/6 has a catalytic role, phosphorylating the transcriptional inhibitor Retinoblastoma protein (Rb) during G1, and partially relieving its inhibition of E2F-mediated transcription. This initiates expression of genes required for cell cycle entry, including Cyclin E. Later in G1, increasing Cyclin E:CDK2 activity results in the hyperphosphorylation and complete inhibition of Rb, allowing full activation of E2F-dependent transcription and entry into S-phase. More recent data has called this model into question, yet still supports a primarily catalytic role for CDK4/6 in cell cycle entry (Narasimha et al., 2014). Indeed, the catalytic activity of CDK4/6 towards Rb has been shown to be a major driver of proliferation(Chung et al., 2019; Harbour et al., 1999; Lundberg & Weinberg, 1998; Topacio et al., 2019; Yang et al., 2020). However, CDK4/6 may also promote cell cycle entry through a non-catalytic role, sequestering the Cip/Kip Cdk inhibitors, p21 and p27, away from CDK2, thus promoting CDK2 activity (Polyak et al., 1994; Sherr & Roberts, 1999). Whilst p21 and p27 inhibit CDK2 activity, they have a more complicated relationship with Cyclin D:CDK4/6. p21 and p27 are necessary for the formation of functional Cyclin D:CDK4/6 complexes and promote complex assembly (Cheng et al., 1999; Guiley et al., 2019; Labaer et al., 1997; Ray et al., 2009). In addition, p27 facilitates the phosphorylation of the T-loop in CDK4 by CDK activating kinase (CAK), which is required for CDK4 kinase activity (Guiley et al., 2019). However, p21/p27 binding can also inhibit Cyclin D:CDK4/6 activity (Guiley et al., 2019; Ray et al., 2009). Other roles for CDK4/6 in cell cycle entry have been suggested (Caillot et al., 2020; Hydbring et al., 2016; Wang et al., 2017). For example, CDK4/6 substrates include proteins controlling mitochondrial function and glycolysis, coordinating the cell cycle and metabolism (Caillot et al., 2020; Salazar-Roa & Malumbres, 2017; Wang et al., 2017). Other studies have reported that CDK4/6 are able to control transcription in a kinase-independent manner (Hydbring et al., 2016; Kollmann et al., 2013). Thus, the precise mechanism by which CDK4/6 activity leads to increased CDK2 activity and cell cycle entry during G1 is unclear.

Our current understanding of CDK4/6 activity suggests two ways by which CDK4/6 inhibitors could act to block proliferation. Our first assumption, based on canonical models of cell cycle entry, is direct CDK4/6 kinase inhibition resulting in cell cycle arrest (Chung et al., 2019; Fry et al., 2004; Toogood et al., 2005). However, it has been reported that CDK4/6 inhibitors are able to arrest cell cycle progression even in the presence of catalytically inactive CDK4/6 (Schade et al., 2019). Further, whilst *RB1* (encoding Rb) status may be an important biomarker for CDK4/6 inhibitor sensitivity, some Rb-deficient tumour cells remain sensitive (Álvarez-Fernández & Malumbres, 2020). An alternative, indirect model of CDK4/6 inhibitor action resolves this issue, implicating CDK2 inhibition as the cause of G1 arrest. Recent experimental work suggests that the CDK4/6 inhibitor Palbociclib can only bind to CDK4 monomers (or potentially also Cyclin D:CDK4 dimers) but not to Cyclin D:CDK4:p21/p27 trimers (Guiley et al., 2019). In this model, cell cycle arrest occurs through inhibition of CDK2 activity by redistribution of p21 and p27 from CDK4 to CDK2 complexes (Guiley et al., 2019). Indeed, resistance to CDK4/6 inhibitors is linked to amplification of Cyclin E and CDK6 which may enable continued proliferation through increased CDK2 activity (Álvarez-Fernández & Malumbres, 2020; Rubin et al., 2020). Increased CDK2 activity has also been reported to result from increased Cyclin D expression, which sequesters p21 and p27 away from CDK2 (Vilgelm et al., 2019). This lack of CDK2 inhibition is proposed to drive proliferation in CDK4/6 inhibitor-treated cells.

CDK4/6 inhibitors are able to arrest cells in G1 despite continued mitogen stimulation (Trotter & Hagan, 2020), indicating that p21 is the most likely candidate for mediating an indirect inhibition of CDK2. Mitogen stimulation results in the abrogation of the inhibitory activity of p27 towards CDK4 due to phosphorylation of the Y74 residue by non-receptor tyrosine kinases (NRTK) such as Src (Chu et al., 2007; Grimmler et al., 2007; Guiley et al., 2019; Hume et al., 2020; Tsytlonok et al., 2019). Further, NRTK signalling can also result in Y88 phosphorylation of p27, which ejects the inhibitory 310 helix of p27 from the CDK2 active site, partially restoring CDK2 activity (Grimmler et al., 2007). Tyrosine phosphorylation of the 310 helix of p21 does not appear to lead to helix ejection, which would allow p21 to retain its function as a CDK inhibitor despite mitogen signalling, which is vital for a robust DNA damage response (Barr et al., 2017; Swadling et al., 2021).

The difference between the two models of CDK4/6 inhibitor action on cell cycle arrest is their dependence on p21 and p27 (Figure 1a). To investigate whether CDK4/6 inhibitors require p21 and p27 to enter or maintain a G1 arrest in cells, we have characterised the expression of p21 and p27 in hTert-RPE1 cells, generated new cell line models to manipulate p21 expression and used live cell imaging to monitor cell cycle arrest in response to Palbociclib. We find that Palbociclib is able to initiate and maintain cell cycle arrest, even when p21 and p27 are removed. Our data call into question the importance of the indirect model of CDK4/6 inhibitor action and suggest that direct inhibition of CDK4/6 by Palbociclib can be sufficient to inhibit proliferation and maintain cell cycle arrest, at least in a non-transformed cellular context.

**Figure 1.**
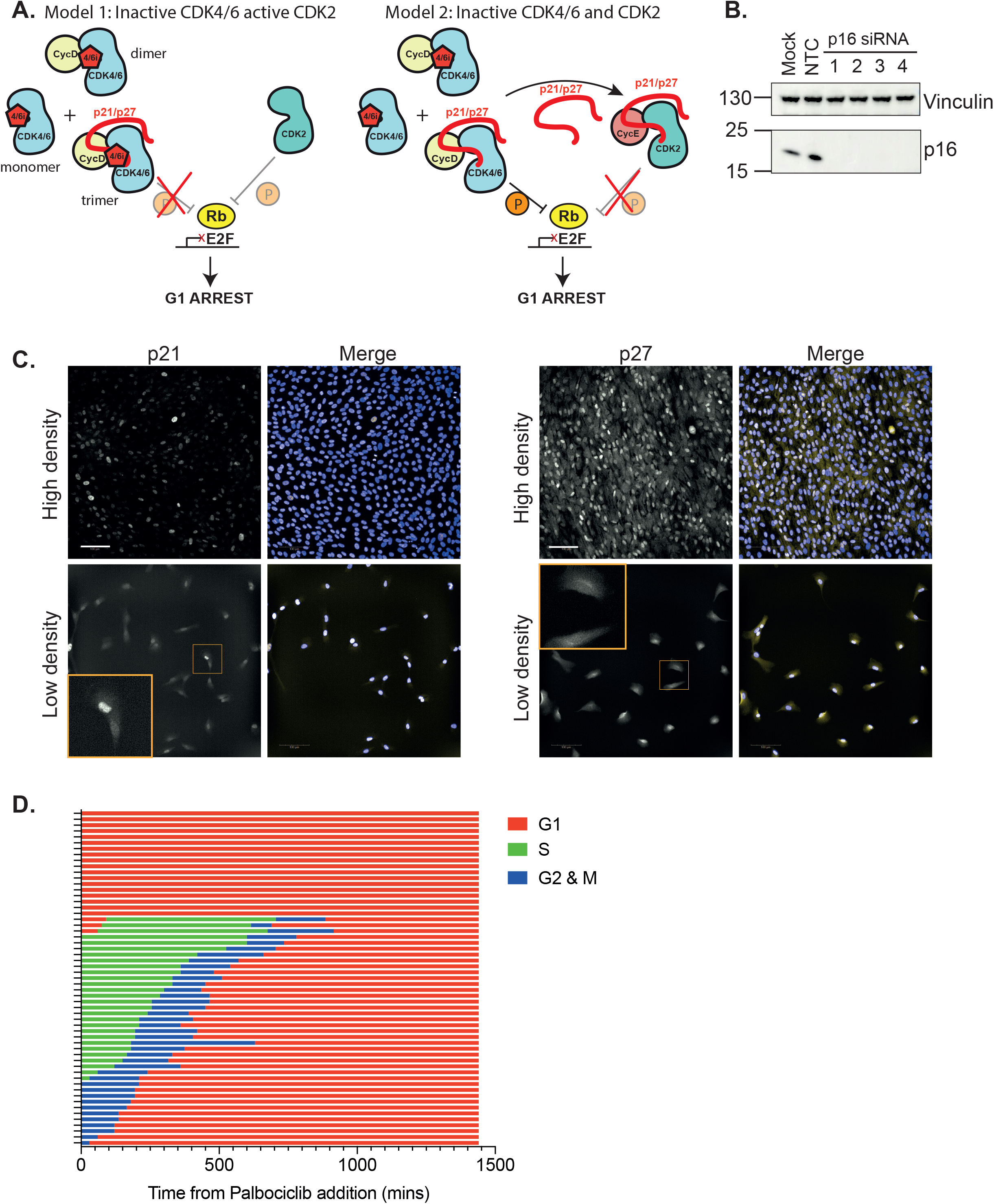
p21 correlates with Cyclin D1 expression and Palbociclib is only effective in G1 in hTert-RPE1 cells. **A.** Models of Palbociclib mechanism of action. Model 1: direct inhibition of CDK4/6 catalytic activity by Palbociclib. Palbociclib binds and inhibits CDK4/6 monomers, CyclinD:CDK4/6 dimers and Cip/Kip:CyclinD:CDK4/6 trimers. Depending when in the cell cycle Palbociclib is added, CDK2 would be active or inactive depending on Cyclin E/A expression. In this model CDK4/6 complexes titrate p21/p27 from CDK2 complexes. Model 2: indirect inhibition of CDK2 activity through redistribution of Cip/Kip inhibitor proteins. Palbociclib binds and inhibits CDK4/6 monomers and CyclinD:CDK4/6 dimers but not Cip/Kip:CyclinD:CDK4/6 trimers. Cip/Kip redistribution from CDK4/6 complexes to CDK2 results in cell cycle arrest. **B.** Western blot showing expression of p16 in hTert-RPE1 cells. Cells were treated with increasing concentrations (1 = 25 nM, 2 = 50 nM, 3 = 75 nM, 4= 100 nM) of siRNA targeting CDKN2A for 48 h. Vinculin in included as a loading control. NTC = Non-targeting control siRNA. **C.** Images show hTert-RPE1 cells plated at high density or low density and fixed and stained for p21 (left-hand side) or p27 (right-hand side; both yellow in merged image) and Hoescht (blue in merged image). Insets in lower panels show magnified images of cells in the image to highlight lack of nuclear p27 in cells cycling at low density. Scale bar is 100 μm. **D.** Graph shows how cell cycle stage affects response to Palbociclib addition. hTert-RPE1 mRuby-PCNA cells were imaged after Palbociclib addition at time 0 and cells were manually analysed to determine cell cycle phenotypes. Each row represents a single cell (n=57 cells) and only one daughter cell was followed post-mitosis. See Supplementary Movie 1 for an example of the imaging data.

## Results

### p21, and not p27, is enriched in the nuclei of cycling hTert-RPE1 cells

For this study we used telomerase-immortalised hTERT-RPE1 cells (RPE1) as they are near diploid, non-transformed, have intact cell cycle control pathways and are sensitive to CDK4/6 inhibitors (Bodnar et al., 1998; Trotter & Hagan, 2020). As such, we assume that cell cycle regulatory complexes will be present at the correct stoichiometries. Previous studies investigating the mechanism of action of CDK4/6 inhibitors have used cancer cells, where the extent to which cell cycle control networks are perturbed is poorly understood. We reasoned that by studying CDK4/6 inhibitor action in RPE1 cells, we could establish a baseline of how Palbociclib modulates a well-controlled cell cycle, which, in the future, can be used to understand the effects of mutations and perturbations observed in cancer cells. We focus on Palbociclib here as it is the best characterised in terms of both its mechanism and its effect on RPE1 cells (Guiley et al., 2019; Trotter & Hagan, 2020).

Whilst RPE1 cells do have reported mutations in CDKN2A and KRAS, there is no clear link between KRAS mutations and Palbociclib sensitivity (di Nicolantonio et al., 2008; Libouban et al., 2017). We confirmed the expression of p16 protein in our RPE1 cells by western blot, indicating that it is not the loss of p16 protein which causes Palbociclib sensitivity in these cells and that cells with functional p16 can still be sensitive (Figure 1b) (Wiedemeyer et al., 2010; Young et al., 2014).

We first characterised the protein expression and localisation of Cip/Kip CDK inhibitor proteins which have been implicated in the response to Palbociclib. Quantification of p21 and p27 levels by immunofluorescence in cycling RPE1 cells, plated at a low density, revealed that p21 protein is exclusively nuclear and is expressed heterogeneously (as previously shown, Barr et al., 2017; Figure 1c). p27 has low nuclear expression in cycling RPE1 cells, and its nuclear levels only increase as cells enter quiescence, as observed when cells are plated at high density and start to enter quiescence through contact inhibition (Figure 1c). This, together with previous data indicating that p21 and Cyclin D levels are correlated in cycling cells (Chen et al., 2013; Labaer et al., 1997; Yang et al., 2017), and that p27 would be largely tyrosine phosphorylated and degraded in growth-factor stimulated cells (Chu et al., 2007; Grimmler et al., 2007; Swadling et al., 2021), suggests that p21, and not p27, is likely to be the primary regulator of Cyclin D:CDK4/6 activity in cycling RPE1 cells. We therefore focussed the majority of our efforts on investigating the role of p21 in the cell cycle response to Palbociclib. However, since the contribution of p27 cannot be discounted, we also perform our assays in the presence and absence of p27.

### Palbociclib is only effective as a cell cycle inhibitor during G1 in RPE1 cells

As it has been established that Palbociclib is limited in its actions to G1 phase (Rubin et al., 2020), we asked when RPE1 cells are sensitive to Palbociclib during the cell cycle with respect to cell cycle arrest. We imaged asynchronous RPE1 cells following Palbociclib addition and followed their cell cycle progression using endogenously tagged mRuby-PCNA (Zerjatke et al., 2017; Supplementary Movie 1). We observed that cells in G1 phase of the cell cycle at the point of drug addition arrest immediately, whilst cells in S, G2 or mitosis complete their current cycle and re-enter G1 phase before arresting (Figure 1d). A small fraction of G1 cells (14.3%) do enter S-phase in the presence of Palbociclib and complete the current cycle before arresting in the next G1. All of these cells enter S-phase within 2 hours of Palbociclib addition and likely represent late G1 cells that were close to the G1/S transition at the time of drug addition. Thus, Palbociclib can only induce cell cycle arrest in cells which are in early/mid G1 phase.

### p21 and p27 are not required for entry into G1 arrest with Palbociclib in RPE1 cells

We hypothesised two possible mechanisms for a G1 arrest response to Palbociclib. One, that consistent with a direct model of Palbociclib action, CDK4/6 activity is only essential for cell cycle progression during early and mid G1 phase of the cell cycle. In this case, whilst CDK4/6 may be inhibited by Palbociclib during the whole cell cycle, this does not affect progression until G1. Alternatively, this could be explained by the indirect model (Figure 1a) as p21 is degraded abruptly at S-phase entry (Barr et al., 2017; Bornstein et al., 2003; Heldt et al., 2018; Nathans et al., 2021) and is therefore only present at high levels during G1 (Rubin et al. 2020).

To test this indirect model of Palbociclib action and determine if cell cycle arrest induced by CDK4/6 inhibitors is dependent upon p21 and p27, we assayed cell cycle distribution by immunofluorescence in fixed cells following 48 h treatment with Palbociclib, in the presence and absence of p21 and p27. We measured EdU incorporation, phospho-S807/811 Rb (P-Rb) levels and DNA content. Cells were pulse labelled with the nucleotide analogue 5-ethynyl-2’-deoxyuridine (EdU) 30 minutes before fixation and Click-iT chemistry used to assay the proportion of cells in S phase (see Methods, Supplementary Figure 2a). Whilst EdU incorporation enabled us to assess the percentage of cells in S phase, P-Rb and Hoechst staining was used to determine cell cycle phase distribution more specifically. Quantification of DNA content by Hoechst sum intensity allows the gating of cells into G1, S and G2/M phases (Supplementary Figure 2b; Chung et al. 2019). P-Rb is bimodally distributed in a population of asynchronously cycling cells, reflecting the proliferation status of the population with G0 cells (and Palbociclib arrested cells) displaying hypophosphorylated Rb (Stallaert et al. 2021, Crozier et al. 2021, Spencer et al. 2013) (Supplementary Figures 2c).

**Figure 2.**
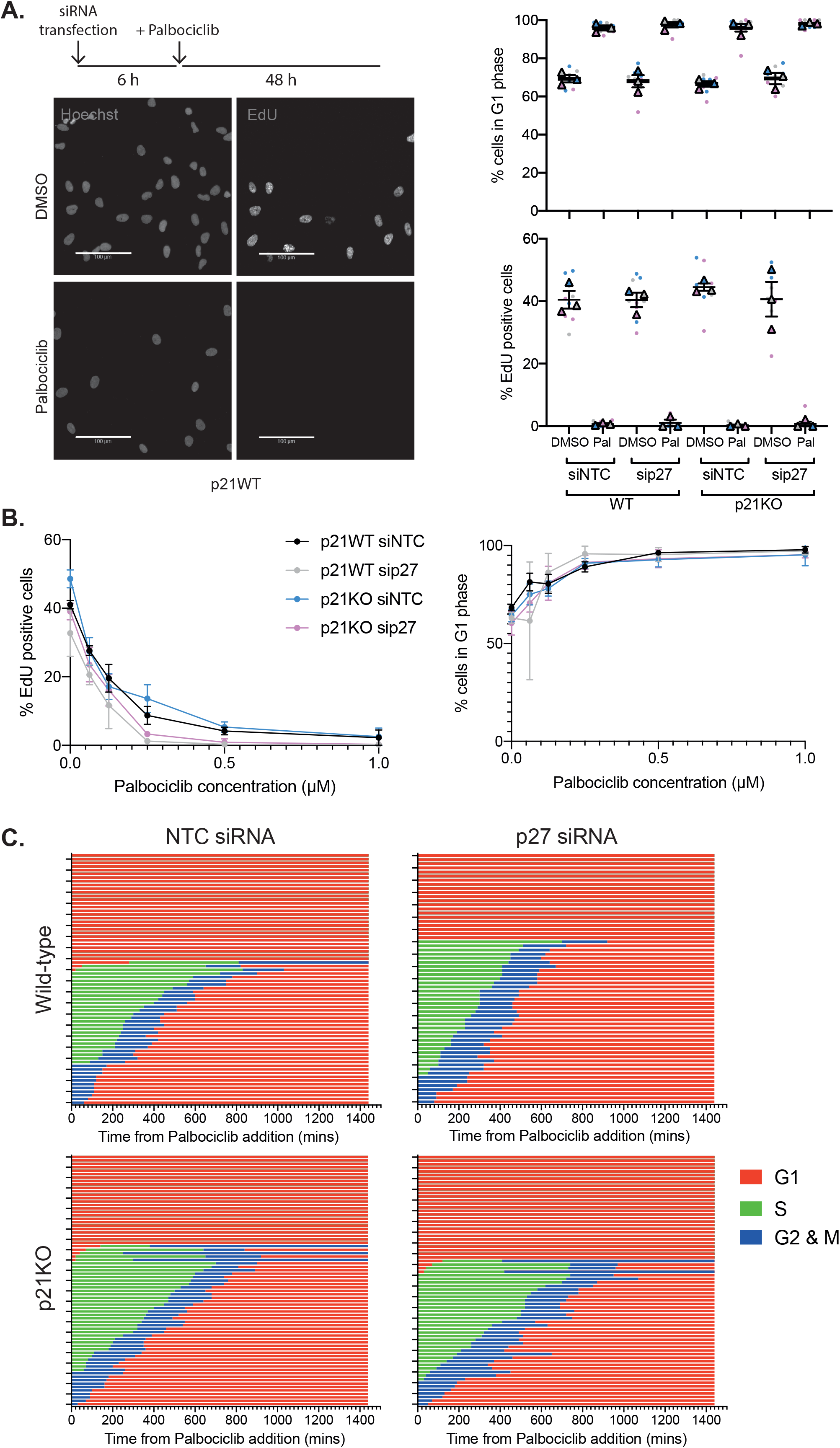
p21 and p27 are not required for Palbociclib-mediated arrest in hTert-RPE1 cells. **A.** hTert-RPE1 p21KO cells were reverse transfected with siNTC or sip27 and treated with DMSO or Palbociclib 6 h after plating, as indicated. Cells were pulse labelled with 10 μM EdU for 30 min before fixation 48 h following drug treatment. EdU positive cells quantified as cells with a nuclear:cytoplasmic ratio of EdU signal as greater than or equal to 1.2. Right, cells were classified in G1 phase according to their DNA content as quantified by Hoechst nuclear sum intensity. Dots in superplots represent replicate wells, colour coded by experimental repeat, triangles represent mean values for each of n=3 experimental repeats with mean and SEM shown. Scale bar, 100 μm. **B.** hTert-RPE1 mRuby-PCNA p21 WT and p21KO cells were treated with the indicated concentrations of Palbociclib for 48 h before 30 min EdU pulse and fixation. Superplot of percentage of EdU positive cells, left, and of cells in G1 phase, right. Mean and SEM from n=3 experimental repeat shown on error bars. **C.** Graphs show timing of cell cycle arrest when Palbociclib is added to asynchronous cells (at 0 mins) and imaged by live cell imaging. Cell cycle phenotypes were monitored and tracked manually over 24 hrs using mRuby-PCNA as a readout. Three fields of view were quantified per condition. Each row represents an individual cell. Wild-type NTC siRNA n=68, wild-type p27 siRNA n=61, p21KO NTC siRNA n=73 and p21KO p27siRNA n=70 cells.

Assessing the cell cycle distribution of untreated p21 knockout (KO) cells (Barr et al., 2017) showed that p21KO does not appreciably alter the fraction of cells in G1 or S phase (Figure 2a columns 1 vs 5), and reduces the fraction of cells with hypophosphorylated Rb compared to p21 wild-type (WT) cells (Supplementary Figure 2d). Assaying the proliferative status of p21KO cells following Palbociclib treatment revealed an arrest in G1 to the same extent as for p21WT cells (Figure 2a columns 2 vs 6, Supplementary Figure 2d).

Whilst we hypothesised that p21 would be more likely than p27 to mediate an indirect mechanism of G1 arrest in Palbociclib in RPE1 cells (Figure 1c), we wanted to ask if Palbociclib could induce arrest in the absence of both CIP/KIP proteins, as p27 has also been implicated in this mechanism (Guiley et al., 2019; Polyak et al., 1994; Sherr & Roberts, 1999). siRNA-mediated knockdown of p27 prior to Palbociclib treatment did not affect proliferation (Figure 2a columns 1 vs 3 and 5 vs 7) or the ability of cells to arrest in G1 (columns 2 vs 4 and 6 vs 8) in p21WT or p21KO backgrounds at a range of Palbociclib concentrations (Figure 2b, Supplementary Figure 2e, f).

Our fixed cell analyses indicated that RPE1 cells arrest in G1 in response to Palbociclib in the absence of p21 and/or p27. However, we wanted to test the hypothesis that it is the presence of p21 (and perhaps p27) during G1 that makes G1 cells sensitive to Palbociclib (Rubin et al., 2020). We reasoned that, if this was the case, then in the absence of p21 and/or p27, a higher fraction of cells in G1 at the time of Palbociclib addition may progress through S-phase and complete the cycle, before arresting in the next G1. Therefore, we repeated our live-imaging experiment in mRuby-PCNA labelled p21WT and p21KO cells treated with non-targeting control (NTC) or p27 targeting siRNA (Barr et al., 2017; Zerjatke et al., 2017). We observed that cells respond in the same way to Palbociclib, arresting in G1, independent of the presence of p21 and p27 (Figure 2c; fraction of G1 cells progressing into S-phase: p21WT NTC 6.5%, p21WT p27si 0%, p21KO NTC 6.7%, p21KO p27si 6.1%).

Together, these data suggest that p21 and p27 are not essential for entry into a Palbociclib-mediated cell cycle arrest and that CDK4/6 activity is only required for cell cycle entry during early and mid G1.

### Generating p21-degron cell lines

In our previous experiments, the absence of p21 and p27 in RPE1 cells at the time of Palbociclib addition means that Palbociclib can bind directly to CDK4 and CDK6 to inhibit their activity (Guiley et al., 2019) and that Palbociclib can only act to arrest the cell cycle through a direct CDK4/6 inhibition mechanism. However, this does not address the question of whether, when present, p21 and p27 are required to mediate an indirect mechanism of cell cycle arrest?

One way to address this question is to allow cells to enter a Palbociclib-mediated arrest in the presence of p21 and p27, and then remove the Cip/Kips and see if any cells re-enter the cell cycle. To be able to efficiently and inducibly degrade p21, we used a double degron system (Hégarat et al., 2020). In this way, we could test if p21 is needed for maintaining a Palbociclib-induced arrest in a system where p21 is normally present to promote the assembly of functional Cyclin D:CDK4/6 complexes, and where p21 could potentially relocalise upon Palbociclib addition to Cyclin:CDK2 complexes to mediate cell cycle arrest (Figure 1a). We introduced an mVenus-mAID-SMASh tag at the C-terminus of p21 in RPE1 cells expressing myc-OsTIR1 under a doxycycline-inducible promoter (p21-degron cells). Homozygous gene targeting was confirmed by PCR and western blot, and an siRNA directed against p21 was used to confirm specificity of tagging (Figure 3a, Supplementary Figure 3a, c). We verified that addition of DIA (doxycycline, IAA and ASV) resulted in the depletion of p21 to undetectable levels by immunoblot after 24 h and that the tag did not affect cell growth and p21 protein localised normally to the nucleus (Figure 3a, Supplementary Figure 3b-d).

**Figure 3.**
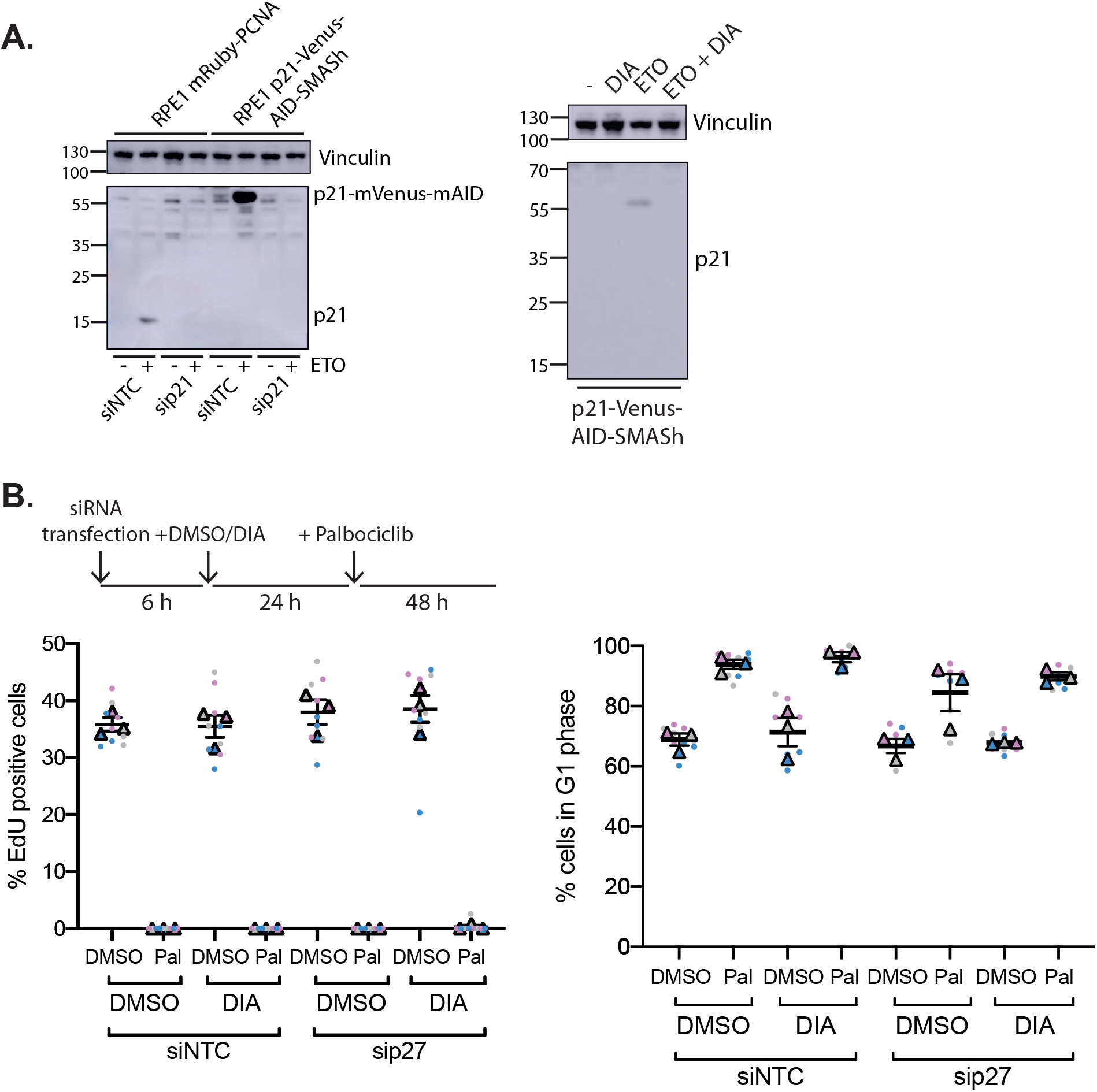
Generation of p21-degron lines **A.** Western blot of whole cell extract from indicated cell lines probing for p21, indicating all p21 expressed in the hTert-RPE1 OsTIR1 mRuby-PCNA p21-mVenus-mAID-SMASh cell line is tagged with mVenus-mAID. Cells were reverse transfected with the indicated siRNAs, non-targeting control (NTC) or p27 and collected after 48 h. 10 μM Etoposide (ETO) was added 24h prior to sample collection to induce DNA damage as p21 expression is low in untreated hTert-RPE1 cells. p21-mVenus-mAID has a predicted molecular weight of 55 kDa, no p21-mVenus-mAID-SMASh is detected as the SMASh tag self-cleaves from the protein. Vinculin was used as a loading control. Right, western blot of whole cell extract indicating that p21-mVenus-AID-SMASh is degraded following DIA (doxycycline, IAA and ASV) addition after induction of p21 expression by ETO. ETO and DIA: Doxycycline (1 μg/ml), IAA (500 μM) and ASV (3 μM) were added 24 h before sample collection. **B.** hTert-RPE1 mRuby-PCNA p21-Venus-AID-SMASh cells were reverse transfected with the indicated siRNAs, 6 h later DMSO or DIA were added as indicated, and 24 h DMSO or Palbociclib added for 48 h. Cells were pulse labelled with EdU before fixation and EdU incorporation and G1 percentage were quantified.

We first used the p21-degron cells to ask if acute depletion of p21 and/or p27 abrogates Palbociclib-induced cell cycle arrest in a system where p21 is normally present to stabilise assembly of Cyclin D:CDK4/6 complexes (in contrast to the p21KO cells which may also have adapted to the loss of p21). Cells were reverse transfected with siRNA, to deplete p27, and treated with DIA, to degrade p21, then treated with Palbociclib 24 h later for 48 h. EdU incorporation, Hoechst and P-Rb staining were used to assess if (and at what cell cycle phase) cells were arrested. Degradation of p21 did not significantly affect the percentage of EdU positive cells, the percentage of G1 phase cells or the distribution of P-Rb staining in DMSO and siNTC treated cells (Figure 3b column 1 vs 3, Supplementary Figure 3e). p21 degradation also did not affect entry into cell cycle arrest mediated by Palbociclib (Figure 3b, column 2 vs 4, Supplementary Figure 3e). Following p27 knockdown alone, proliferation was largely unaffected, similar to what we observed in p21WT and p21KO cells (Figure 3b, columns 1 vs 5, Supplementary Figure 3e). p27 knockdown also did not affect entry into cell cycle arrest in Palbociclib, independently of DIA addition prior to treatment (Figure 3b columns 2 vs 6 and 4 vs 8, Supplementary Figure 3e).

In summary, neither p21 nor p27 are necessary for the initiation of a Palbociclib-mediated arrest in RPE1 cells.

### p21 and p27 are not required for maintenance of G1 arrest with Palbociclib

To further clarify the importance of direct or indirect mechanisms of Palbociclib action in our system, we wanted to ask if maintenance of cell cycle arrest initiated in unperturbed conditions is dependent on p21 and p27. We reasoned that if an indirect mechanism maintains cell cycle arrest during Palbociclib treatment then a decrease in p21 or p27 protein levels during arrest would result in cell cycle re-entry.

To test if removal of p21 and/or p27 promoted cell cycle re-entry in Palbociclib-arrested cells, we first degraded p21 following 24 h Palbociclib treatment in p21-degron cells. Assaying proliferation as before, we saw that cell cycle arrest was maintained following p21 degradation (Figure 4a columns 5 vs 6, Supplementary Figure 4a). Further, p27 knockdown following Palbociclib treatment did not affect the arrest, independent of the presence of p21 (Figure 4a columns 7 vs 8, Supplementary Figure 4a). Additionally, in Palbociclib treated p21KO cells, the knockdown of p27 did not affect the arrest (Figure 4b columns 7 vs 8, Supplementary Figure 4b).

**Figure 4.**
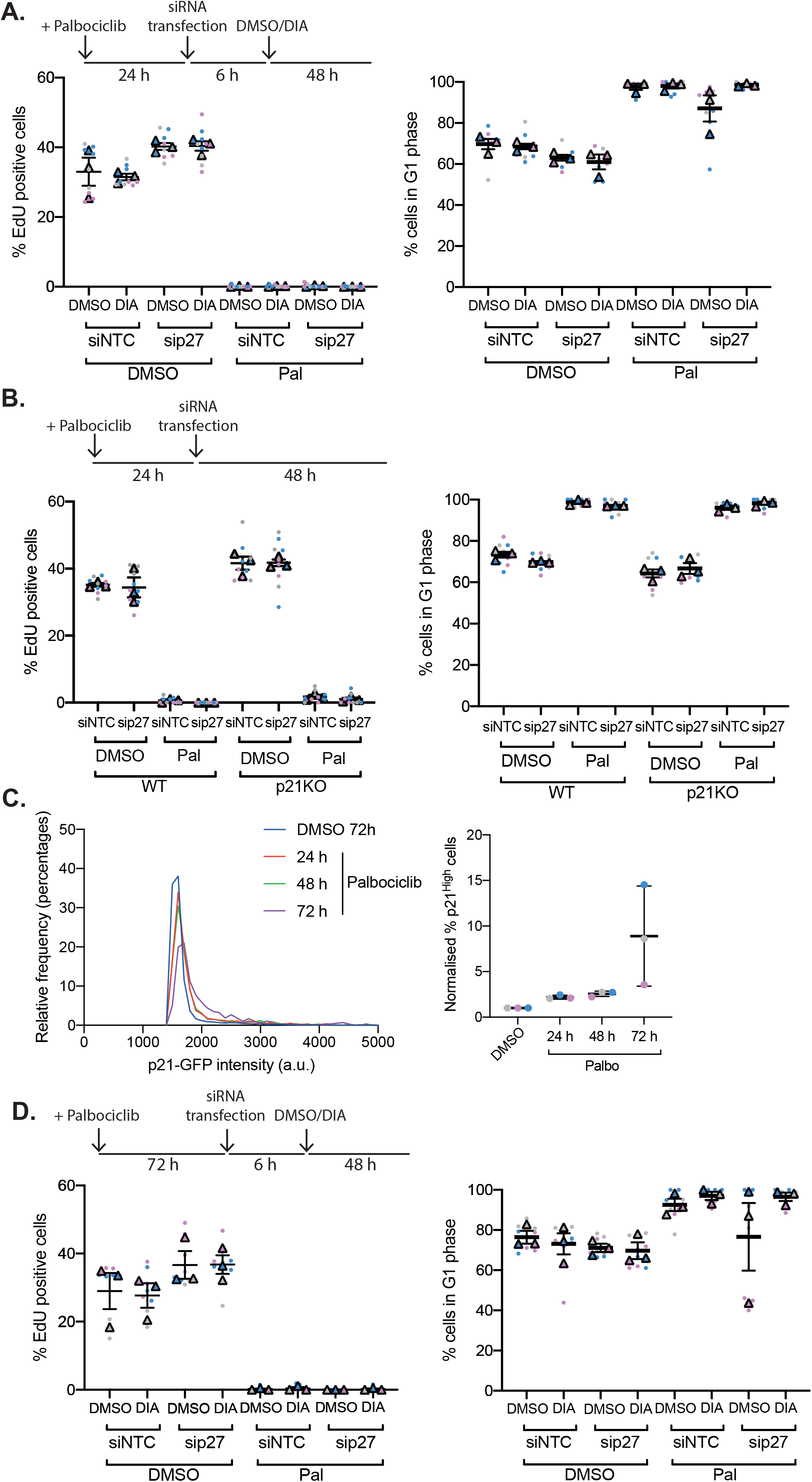
Palbociclib-dependent arrest can be maintained in the absence of p21/p27. **A.** hTert-RPE1 p21WT and KO cells were treated with DMSO/Palbociclib, transfected with the indicated siRNAs 18 h later, then treated with DIA 6 h following transfection. Cells were pulse labelled for 30 minutes with 10 μM EdU and fixed 48 h following DIA addition. Dots represent replicate wells, colour coded by experimental repeat. Triangles represent mean values for each experimental repeat with mean and SEM shown. **B.** hTert-RPE1 p21-degron cells were treated with DMSO/Palbociclib, transfected with the indicted siRNAs 24 h after drug treatment, then treated with DMSO/DIA 6 h following transfection. 48 h after DIA addition, cells were pulse labelled with 10 uM EdU for 30 minutes and fixed. **C.** hTert-RPE1 mRuby-PCNA p21GFP cells were treated with DMSO for 72 h or Palbociclib for the indicated times, and p21GFP levels quantified. Representative frequency distribution of measured intensities from one experimental repeat shown. Right, percentage of cells classified as p21 high above a threshold of p21 intensity, n=3 experiments shown. Data was normalised to 72 h DMSO treatment within each experimental repeat. **D.** hTert-RPE1 p21-degron cells were treated with DMSO/Palbocilib for 72 h, transfected, then DMSO/DIA added 6 h later. Cells were pulse labelled with 10 μM EdU for 30 minutes 48 h after DIA addition.

We observed that Palbociclib treatment induces an increase in p21 and p27 protein levels in cells in a time-dependent manner, and that p21 localisation remains exclusively nuclear. The largest increase in p21 protein occurs between 48 and 72 h Palbociclib treatment (p21: Figure 4c; p27: Supplementary Figure 2d columns 1 vs 2). We hypothesised that this might reflect an increased dependence on p21 and p27 to maintain cell cycle arrest in the presence of Palbociclib. To test this, we decreased p21 and p27 protein levels following a long-term Palbociclib-mediated arrest, to ask if these proteins are necessary to maintain a prolonged arrest initiated in unperturbed conditions. We used p21-degron cells to degrade p21 72 h following Palbociclib treatment. Assaying proliferation 48 h following the induction of p21 degradation revealed a similar extent of arrest, independent of the presence of p21 (Figure 4d columns 5 vs 6, Supplementary Figure 4c) or p27 (columns 5 vs 7 and 7 vs 8).

Together, our data demonstrate that RPE1 cells are not dependent on p21 or p27 for maintenance of a Palbociclib-mediated cell cycle arrest.

## Discussion

In this study, we have established that a Palbociclib-mediated cell cycle arrest is not dependent on the CIP/KIP inhibitor proteins p21 and p27 in RPE1 cells. Using this non-transformed cell line, we have demonstrated that cell cycle arrest in response to Palbociclib can both be initiated and maintained without p21 or p27.

Importantly, in a system in which a ‘normal’ Palbociclib-mediated arrest has been allowed to occur, the presence of p21 and p27 is not necessary for the maintenance of cell cycle arrest (Figure 4). This indicates that an arrest initiated in the presence of Cyclin D:CDK4/6 trimeric complexes with p21 (and potentially p27), which might be predicted to occur through an indirect mechanism of action of Palbociclib, is not dependent on p21/p27. If the arrest were maintained through the indirect inhibition of CDK2, we would predict that the absence of p21/p27 would result in the release of, at least a fraction of, cells into the cell cycle despite the continued presence of Palbociclib. Whilst we are unable to rule out that in the presence of p21 and p27, Palbociclib acts through an indirect mechanism to inhibit CDK2 to initiate arrest that is then maintained by direct Palbociclib-mediated CDK4/6 inhibition, our data suggests that the presence of p21 and/or p27 is not essential for entry into cell cycle arrest or maintenance of that arrest in a non-transformed cell line.

This calls into question a solely indirect model of Palbociclib driven cell cycle arrest, which is dependent upon the presence of p21 and/or p27 both before and during the arrest (model 2, Fig 1A). Our data support the direct inhibition of CDK4/6 by Palbociclib to inhibit proliferation. This is supported by work assaying CDK4/6 and CDK2 activity in single non-transformed MCF10A cells using live cell CDK4/6 and CDK2 activity reporters (Yang et al., 2020). In both G1 and S phase cells released from synchronisation in G0, Palbociclib addition decreases CDK4/6 activity within one hour, while CDK2 activity decreases at a much slower rate. Further, recent data from multiple cell line models suggested that in contrast to CDK2, CDK4 catalytic activity towards Rb is inhibited by Palbociclib treatment (Simoneschi et al., 2021). Together, this suggests that Palbociclib directly inhibits CDK4/6 catalytic activity and that this is sufficient for a G1 phase arrest.

Whilst sensitivity to Palbociclib is known to be restricted to G1, here we have reported that cells become insensitive to drug addition in late G1, at approximately two hours before S phase entry (Figure 1d). This corresponds with early reports of restriction point timing (Campisi et al., 1982; Foster et al., 2010; Yen & Pardee, 1978). This could reflect an increasing rate of p21 degradation as cells approach the G1/S transition (Heldt et al., 2018; Nathans et al., 2021) or could be the result of a change in the dependency of cells on CDK4/6 activity for cell cycle progression at the restriction point. Since we see no change in sensitivity of G1 cells to Palbociclib in the absence of p21 and/or p27, it is likely that it is the latter hypothesis that is correct here and that cells only require CDK4/6 activity in early and mid G1 to complete the cell cycle.

Interestingly, p21 has been implicated in cellular resistance mechanisms to CDK4/6 inhibitors, indicating it still has an important role in their mechanism of action in some contexts. The loss of p53, a major driver of p21 expression, has been implicated in resistance to the CDK4/6 inhibitor Abemaciclib, with a significant enrichment in *TP53* mutations in resistant breast cancer (Patnaik et al., 2016; Wander et al., 2020). Further, increasing p21 expression is linked to re-sensitising resistant cells to Palbociclib, indicating that low p21 levels may contribute to Palbociclib resistance (AbuHammad et al., 2019; Vilgelm et al., 2019). However, loss of p53 does not prevent proliferative arrest induced by CDK4/6 inhibitors, supporting our hypothesis that CDK4/6 inhibitors are able to act through multiple potential mechanisms (Wander et al., 2020). In contrast, Y88 phosphorylation of p27, a modification which prevents its inhibitory activity towards CDK2, correlates with sensitivity to Palbociclib (Gottesman et al., 2019).

Our data does not rule out a potential role for p21/p27 during Palbociclib-induced cell cycle arrest in some cells (Guiley et al., 2019). Indeed, it indicates that Palbociclib is able to arrest the cell cycle through parallel direct and indirect mechanisms and that the dominant mechanism depends upon the cellular context. A parallel pathways model explains how both RPE1 cells acutely depleted of p21/p27 and cells with impaired CDK4/6 or Rb activity are sensitive to Palbociclib (Guiley et al., 2019; Schade et al., 2019; Zhao & Burgess, 2019).

This could represent a difference between healthy and cancer cells in their dependence on CIP/KIP proteins for arrest. Whilst MCF7 breast cancer cells are at least partly dependent on CDK4/6 activity for cell cycle entry (Grillo et al., 2006), p21/p27 appear to mediate their Palbociclib sensitivity (Guiley et al., 2019). It therefore seems likely that Palbociclib is able to arrest cell cycle progression through both direct and indirect mechanisms, meaning the sensitivity of a cancer cell to Palbociclib may be dependent upon both its reliance on CDK4/6 activity for cell cycle entry and the relative expression levels of p21/p27. As the activity of these pathways is often perturbed in cancer, this may alter the effect of Palbociclib on the cell cycle, determining a cell’s sensitivity to Palbociclib and the mechanism by which it may cause cell cycle arrest. For example, the lack of sensitivity of some triple-negative breast cancer cells (TNBC) may reflect both a decreased dependence on CDK4/6 activity for cell cycle entry (due to high Cyclin E expression and CDK2 activity) and low p21 and/or p27 levels (Asghar et al., 2017). The prediction of a cell’s sensitivity to Palbociclib may therefore require information about the balance between the activity of multiple cell cycle pathways (Table 1). For example, we would predict that in Rb-deficient cells which remain sensitive to Palbociclib would be sensitive to decreases in p21/p27. These different potential mechanisms of action of Palbociclib may explain why there are no clear biomarkers for sensitivity.

**Table 1.** Predictions of sensitivity to CDK4/6 inhibitors.

| Dependence on CDK4/6 | p21/p27 levels | CDK4/6 inhibitor sensitivity prediction |
| --- | --- | --- |
| Yes | High/normal | Yes – both mechanisms |
| Yes | Low | Yes – direct inhibition |
| No | High | Yes – indirect mechanism |
| No | Low | No |

## Authors’ contributions

A.R.B. conceived the study. A.R.B and B.R.P. designed, performed and analysed the experiments and wrote the manuscript.

## Acknowledgements

We would like to thank Adrian Saurin and Tony Ly for critical reading and discussion of the manuscript. We also thank the LMS/NIHR Imperial Biomedical Research Centre Flow Cytometry Facility for the support. BRP and ARB are funded by CRUK CDF: C63833/A25729 and MRC core funding to the London Institute of Medical Sciences (MC-A658-5TY60).

## Methods

### Cell culture

hTert-RPE1 cells were from ATCC and were maintained in DMEM (Gibco) supplemented with 10% FBS and 1% Penicillin-Streptomycin at 37°C and 5% CO2. RPE1 mRuby-PCNA p21-GFP cells, in which both alleles of the endogenous CDKN1A locus were labelled with GFP at the C-terminus and one allele of PCNA was labelled at the N-Terminus with mRuby, were described previously (Barr et al., 2017). RPE1 mRuby-PCNA p21 KO 1A cells were described previously (Barr et al., 2017).

Drugs used and working concentrations: Etoposide 10 μM, Doxycycline 1 μg/ml, Indole-3-acetic acid (IAA) 500 μM, Asunaprevir (ASV) 3 μM, Palbociclib 1 μM (unless otherwise stated).

### Generation of p21-Venus-AID-SMASh tagged hTert-RPE1 cell line

An mVenus-mAID-SMASh tag was introduced to the C terminus of the human CDKN1A gene using targeting vectors and gRNA/Cas9 cleavage.

For the homology donor plasmid primers used for the left and right homology arms were the same as in Barr et al. 2017. To PCR amplify mVenus, we used the following primers: forward, 5’-TCTTCTCCAAGAGGAAGCCCGGAGGAGGAGTGAGCAAGGGCGAGGAG-3’, reverse 5’-GCTGATGCCGCTGAGGCGCCCTTGTACAGCTCGTCCAT-3’. mAID-

SMASh-Neomycin was amplified with the primers forward: 5’-GGCGCCTCAGCGGCATCAGCTGCAGGAGCTGGAGGTGCATC-3’ and reverse: 5’-GCAGGCTTCCTGTGGGCGGATCAGAAGAACTCGTCAAGAAG-3’. LHA, mVenus, mAID-SMASh-Neomycin, RHA PCR products were ligated into pAAV p21 vector by Gibson assembly at a ratio vector:inserts of 1:2:2 using T4 DNA ligase (NEB). All constructs were checked by sequencing before transfection into cells. To generate stable clones, hTERT-RPE1 OsTIR1 cells (a gift from Helfrid Hoechegger, Hégarat et al., 2020) were transfected with pX330 g21 gRNA plasmid (Barr et al., 2017) and the p21 homology donor plasmid at a ratio of 1:1 using Lipofectamine 2000, according to the manufacturer’s instructions (Invitrogen). Cells were incubated for 3 weeks in media containing 0.5 μg/ml G418 and selected clones were screened by western blot and genomic DNA PCR.

### siRNA transfection

Cells were transfected with siRNA at a final concentration of 20 nM using Lipofectamine RNAiMAX, according to the manufacturer’s instructions (Invitrogen). Briefly, 40 nl of Lipofectamine RNAiMAX (Invitrogen) was mixed with siRNA in 10 μl OptiMEM (Gibco) per well of a 384 well plate. 20 μl of cells at a density of 2.5×10^4^ cells/ml were plated on top of this, and cells were incubated at 37 °C. siRNAs used were Dharmacon ON-TARGETplus Non-targeting siRNA #1 (NTC) and CDKN1A (set of 4), Ambion Silencer Select siRNA CDKN1B (Cat. No 4427038) and p16 siRNA sequence used: UACCGUAAAUGUCCAUUUAUA.

### Immunofluorescence

Cells were grown on 384 well CellCarrierUltra (PerkinElmer) plates. For EdU staining, a final concentration of 10 μM EdU was added to the growth media 30 minutes prior to fixation. Cells were fixed in 4% paraformaldehyde in PBS for 15 minutes, washed three times with PBS. Permeabilization in PBS 0.2% Triton X-100 for 15 minutes was followed by blocking in 2% BSA in PBS for 1 hour. Cells were incubated with primary antibodies diluted in blocking buffer at 4°C overnight, washed three times with PBS then incubated with a 1:1000 dilution of secondary antibodies for 1 hour at room temperature. For EdU detection cells were incubated for 30 minutes in TBS 100 mM pH 7.5, CuSO_4_ 4mM, Sulfo-cyanine 3 azide 5 μM, sodium ascorbate 100 mM. Cells were washed three times in PBS, incubated for 10 minutes with 1 μg/ml Hoechst, then washed a further three times in PBS.

Antibodies used: p21 (Invitrogen MA5-14949 1:1000), p27 (CST 3688 1:1000), P-Rb S807/811 (CST 8516, 1:2000); secondary Goat anti-rabbit IgG (H+L) Alexa Fluor 647 (Invitrogen A21245, 1:1000). Plates were imaged using a 20X (NA 0.8) objective using an Operetta CLS microscope.

### Western blot

Whole cell extract of RPE1 cells was collected following aspiration of medium from culture plate, two washed in PBS and the addition of 1X Novex Tris-glycine SDS sample buffer (Invitrogen) and collection of cells by scraping. Samples were incubated at 95°C for 10 minutes before loading on 12-15% precast NuPAGE gels (Invitrogen).

Primary antibodies used: p16 (CST 80772, 1:1000), p21 (Invitrogen MA5-14949 1:1000), vinculin (CST 13901 1:1000); secondary antibodies HRP linked Anti Rabbit IgG (CST 7074, 1:2000).

### Growth curves

Cells were plated at a density of 20,000 cells per well in duplicate in 6-well plates. Brightfield images were taken every 2 hours for 5 days and the percentage confluency calculated using an Incucyte Live-Cell analysis system (Sartorius).

### Live imaging of Palbociclib addition

hTert-RPE1 cells were seeded into 384 well CellCarrier Ultra plates (PerkinElmer) one day prior to imaging at a density of 1000 cells/well in 20 μl of phenol-red free DMEM:F12 with 10% FBS and 1% P/S. In cases where cells were transfected with siRNA, cells were plated onto siRNA:lipofectamine RNAiMax (Invitrogen) complexes (as described elsewhere). Prior to imaging, media was added to all wells to a final volume of 100 μl, with a final concentration of 1 μM Palbociclib. A breathable film was applied to the plate (ThermoFisher) to prevent media evaporation and cells were imaged on the Operetta CLS (PerkinElmer) at 37 °C and 5% CO2, using a 20x (N.A. 0.8) objective, every 10 (Figure 2) or 15 (Figure 1) mins. Image analysis was performed in FIJI and NucliTrack (Cooper et al., 2017). Endogenously tagged mRuby-PCNA was used, as previously described to determine cell cycle timing (Zerjatke et al., 2017).

