## Supplementary Information for "Palbociclib-mediated cell cycle arrest can occur in the absence of the CDK inhibitors p21 and p27"

A.

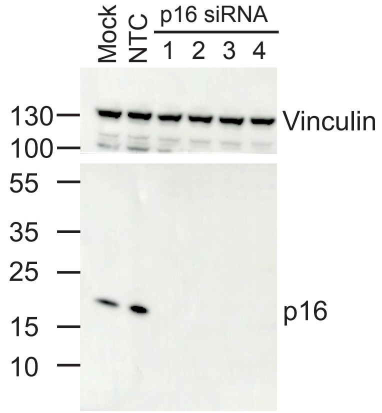

**Supplementary Figure 1. A.** Uncropped western blot shown in Figure 1b.

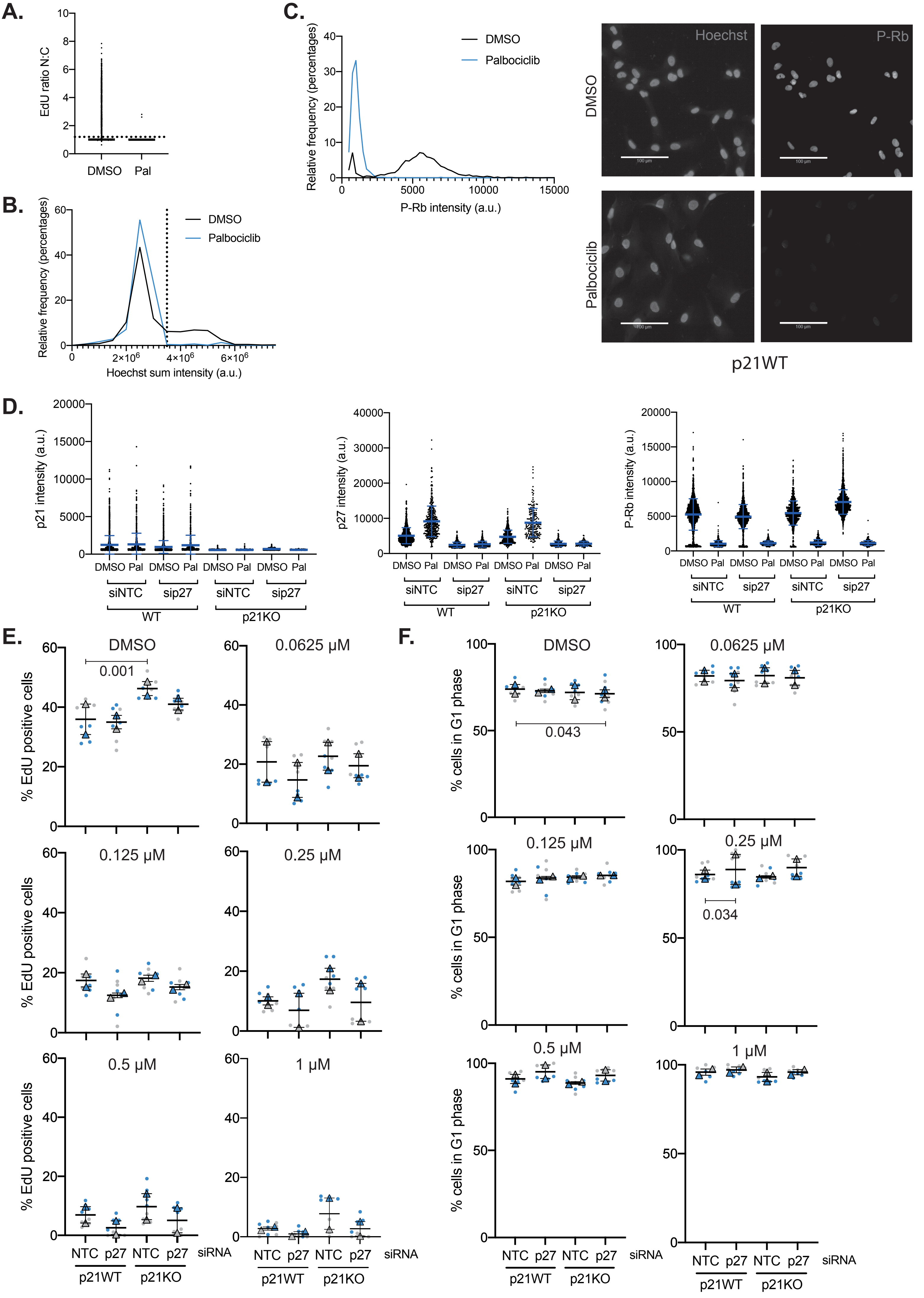


**Supplementary Figure 2**

**Supplementary Figure 2. A-C** Rationalisation of methods for quantification of cell proliferation. hTert-RPE1 mRuby-PCNA p21WT were treated as in Figure 2a, only data for siNTC and DMSO treated cells shown for representative of n=3 experimental repeats. **A.** EdU positive cells were quantified as having a nuclear:cytoplasmic ratio of EdU signal above 1.2, indicated by the dotted line. **B.** Cells were quantified as being in G1 phase if their Hoechst sum intensity was below a threshold level decided by Hoechst sum intensity distribution in p21WT siNTC DMSO treated cells. Threshold for this experimental repeat shown as dotted line. Data for p21WT siNTC Palbociclib treated cells overlaid for comparison. **C.** Frequency distribution of P-Rb levels in individual nuclei of p21WT siNTC DMSO/Palbociclib treated cells with representative images, scale bar 100 μm **D.** hTert-RPE1 mRuby-PCNA p21WT and p21KO were treated as in Figure 2a. Protein levels of p21, p27 and P-Rb were quantified by immunofluorescence. **E, F** hTert-RPE1 mRubyPCNA p21 WT and p21KO cells were treated as in Figure 2b. Superplots of data from n=3 experimental repeats showing percentage of EdU positive cells (**E**) and G1 cells (**F**) at different concentrations of Palbociclib or DMSO. Significance was calculated within each concentration using a non-parametric Kruskal-Wallis test with Dunn’s multiple comparisons test comparing each condition to p21WT siNTC treated cells. Only p values <0.05 are shown, all other comparisons were not significant.

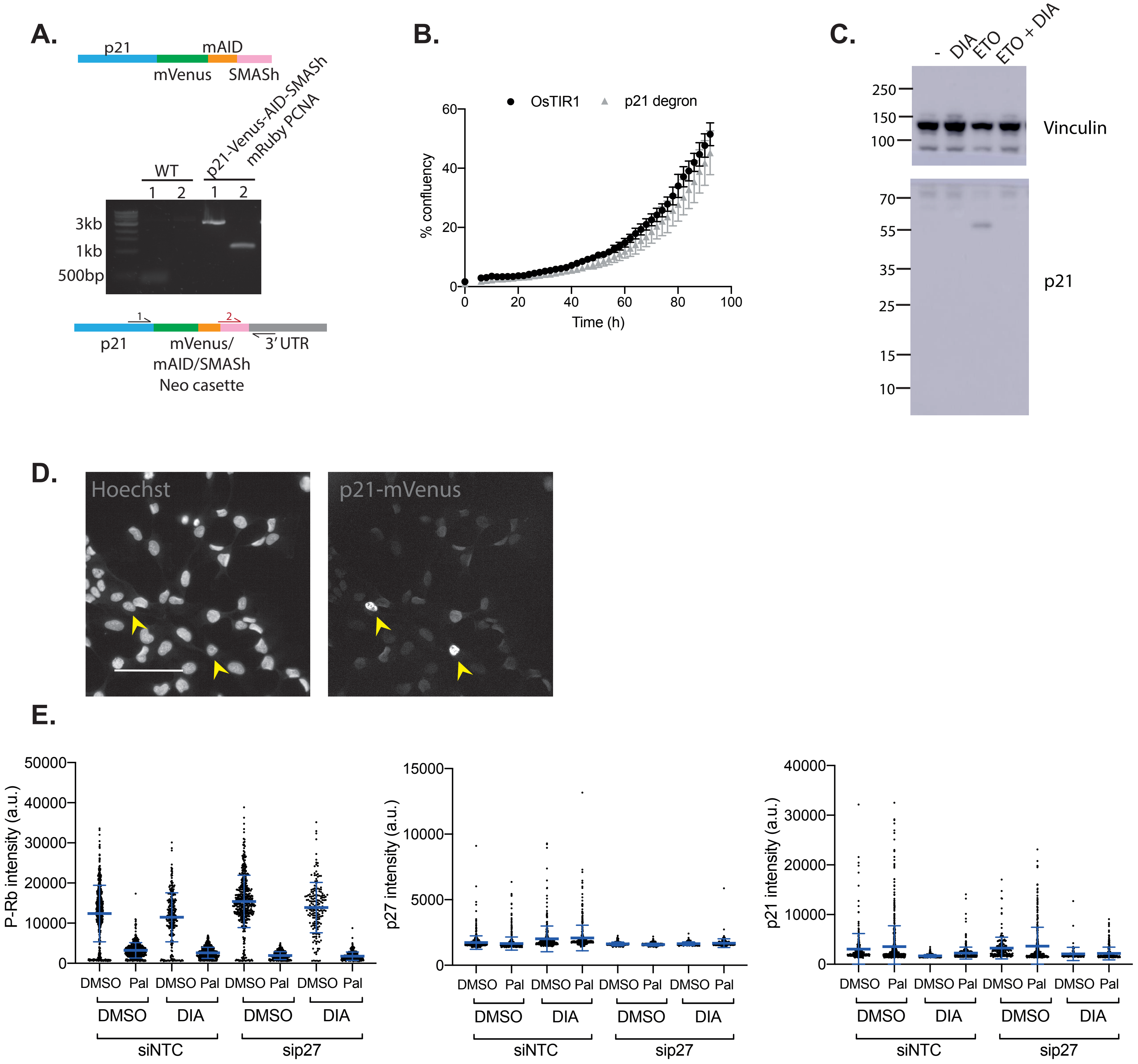


**Supplementary Figure 3**

**Supplementary Figure 3. A.** Schematic of p21-Venus-AID-SMASh. Genomic DNA was prepared from hTert-RPE1 mRuby-PCNA cells with wild type p21 and p21 endogenously tagged with mVenus-AID-SMASh. PCR was performed on the genomic DNA using two different forward primers, 1 and 2, as indicated. Expected band size with; forward primer 1: WT cells 477 bp, p21-degron cells 2 kbp; primer 2: WT no band, p21-degron cells 1048 bp. **B.** Growth curve of hTert-RPE1 OsTIR1 and hTert-RPE1 OsTIR1 p21-mVenus-AID-SMASh cells. Cells were imaged every 2 hours for 5 days and the mean and SD confluency from two wells plotted. Calculated as the percentage of total well area covered. **C.** Uncropped western blot shown in Figure 3a, right. **D.** Representative image of hTert-RPE1 p21-Venus-AID-SMASh cells, scale bar 100 μm. Nucleus with high p21 levels marked with arrowhead. **E.** hTert-RPE1 p21-degron cells were treated as in Figure 3b. Representative graphs, from n=3 experiments, of quantification of P-Rb, p27 and p21 levels by immunofluorescence. Each dot represents one cell, mean and SD are shown in blue.

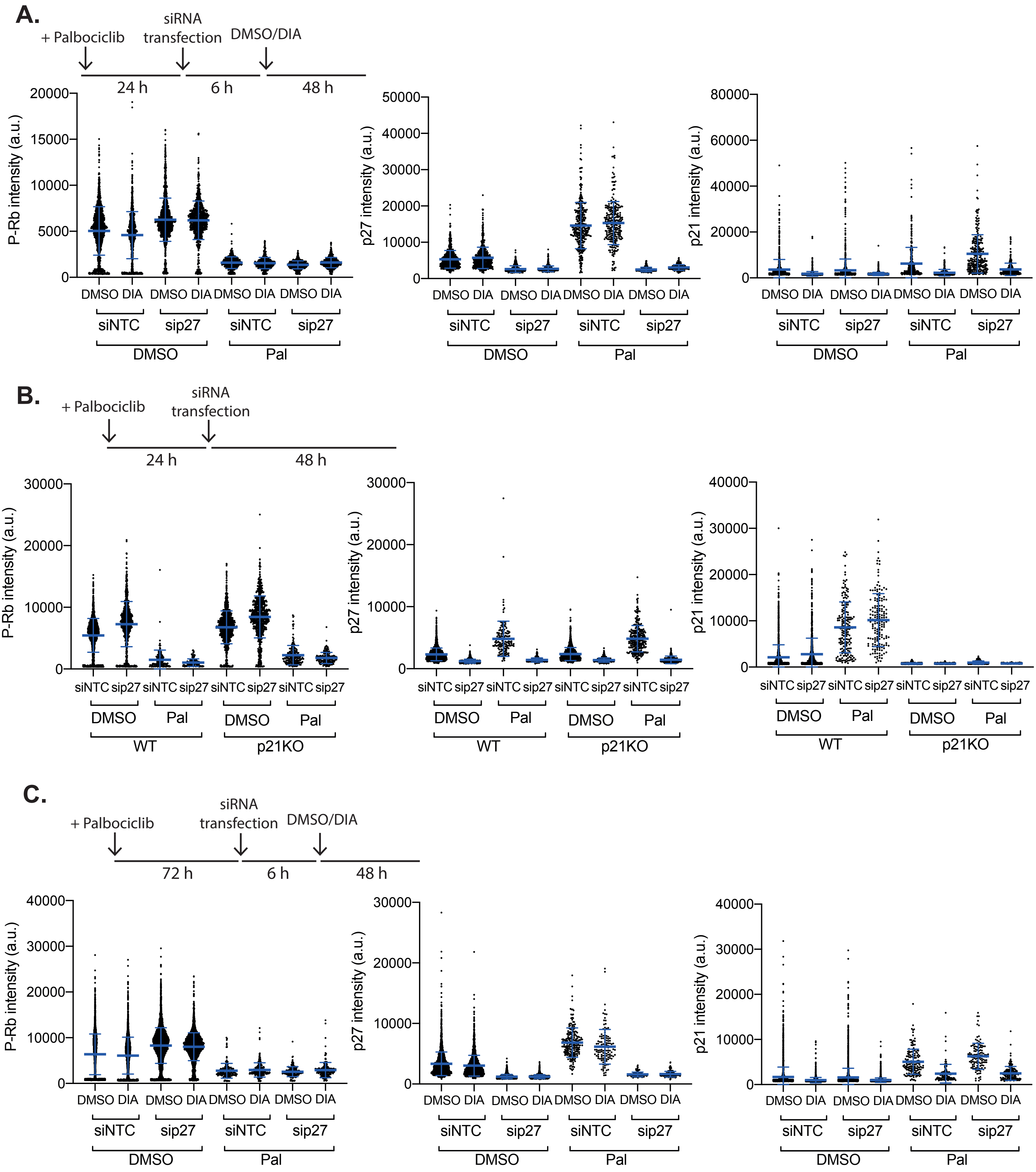


**Supplementary Figure 4**

**Supplementary Figure 4. A-C.** hTert-RPE1 p21-degron cells were treated as in Figure 4a, b and d, respectively. P-Rb, p21 and p27 protein levels were quantified by immunofluorescence. Representative plots of single cell data shown from n=3 experimental repeats.

**Supplementary Movie Legends**

**Supplementary Movie 1.** hTert-RPE1 mRuby-PCNA cells treated with 1$\mu$M Palbociclib at time 00:00. Images were captured every 15 min. Loss of mRuby-PCNA signal indicates entry into G0/G1 arrest.
